# Soft Substrate Maintains Proliferative and Adipogenic Differentiation Potential of human Mesenchymal Stem Cells on Long Term Expansion by Delaying Senescence

**DOI:** 10.1101/364059

**Authors:** Sanjay K. Kureel, Pankaj Mogha, Akshada Khadpekar, Vardhman Kumar, Rohit Joshi, Siddhartha Das, Jayesh Bellare, Abhijit Majumder

## Abstract

Human mesenchymal stem cells (hMSCs), when cultured on tissue culture plate (TCP) for *in vitro* expansion, they spontaneously lose their proliferative capacity and multi-lineage differentiation potential. They also lose their distinct spindle morphology and become large and flat. After a certain number of population doubling, they enter into permanent cell cycle arrest, called senescence. This is a major roadblock for clinical use of hMSCs which demands large number of cells. A cell culture system is needed which can maintain the stemness of hMSCs over long term passages yet simple to use. In this study, we explore the role of substrate rigidity in maintaining stemness. hMSCs were serially passaged on TCP and 5 kPa poly-acrylamide gel for 20 population doubling. It was found that while on TCP, cell growth reached a plateau at cumulative population doubling (CPD) = 12.5, on 5 kPa gel, they continue to proliferate linearly till we monitored (CPD = 20). We also found that while on TCP, late passage MSCs lost their adipogenic potential, the same was maintained on soft gel. Cell surface markers related to MSCs were also unaltered. We demonstrated that this maintenance of stemness was correlated with delay in onset of senescence, which was confirmed by β-gal assay and by differential expression of vimentin, Lamin A and Lamin B. As preparation of poly-acrylamide gel is a simple, well established, and well standardized protocol, we believe that this system of cell expansion will be useful in therapeutic and research applications of hMSCs.

**One Sentence Summary:** hMSCs retain their stemness when expanded *in vitro* on soft polyacrylamide gel coated with collagen by delaying senescence.

**Significance Statement:** For clinical applications, mesenchymal stem cells (MSCs) are required in large numbers. As MSCs are available only in scarcity *in vivo*, to fulfill the need, extensive *in vitro* expansion is unavoidable. However, on expansion, they lose their replicative and multi-lineage differentiation potential and become senescent. A culture system that can maintain MSC stemness on long-term expansion, without compromising the stemness, is need of the hour. In this paper, we identified polyacrylamide (PAA) hydrogel of optimum stiffness that can be used to maintain stemness of MSCs during *in vitro* long term culture. Large quantity of MSCs thus grown can be used in regenerative medicine, cell therapy, and in treatment of inflammatory diseases.

## Introduction

Human Mesenchymal Stem Cells (hMSCs), due to their multi-lineage differentiation potential, immuno-suppressive capability, and immuno-modulatory effects, have been used with varying degree of success to treat cardiovascular, musculoskeletal, immune related and haemopoietic diseases (*1–4*). Though MSCs are available from multiple adult tissues (5), critically low availability of MSCs in isolated sample is a major roadblock for clinical trials. For example, in bone marrow aspirates, only 0.001 to 0.01% of the nucleated cells are MSCs (6) while, a dose of roughly about 100 million cells are required to treat a person of 70 kg of weight (7). As a result, a long term *in vitro* expansion is essential to reach a significant number of cells for autologous treatment.

However, on *in vitro* expansion, MSCs lose their proliferative ability and multilineage differentiation potential (8,9). Like any other primary somatic cells, after certain number of cell divisions, they enter senescence state which is morphologically characterized by enhanced spreading area and shape irregularity (*10,11*). More importantly, they lose their multilineage potential, migration and homing ability (*12,13*) making them unsuitable for clinical use (*14,15*). Though multiple approaches have been tried to maintain MSC stemness over prolonged expansion (*16*), finding a culture system to achieve the same is still an unmet need. In this context, it might be of interest to mention that NIH in their website has listed six points that are needed to be addressed to realize the potential of stem cell-based therapies. The first one in that list is “Stem cells must be reproducibly made to proliferate extensively and generate sufficient quantities of cells for making tissue” (17). A culture system that can fulfill this need may help to progress regenerative medicine significantly.

Controlling physical microenvironment of our cell culture system might offer a solution in this context. In past 15 years, it has been shown that mechanical cues such as stiffness of cell culture substrate, shear stress, mechanical strain, cell morphology, substrate topology etc. influence wide array of cell behavior and cell fate including survival, proliferation, and differentiation (*18–24*). It has also been shown that such mechanical cues may play important role in maintaining MSCs stemness. For example, MSCs cultured on micro-contact printed islands, as spheroids, and on nanopatterns were shown to retain multipotency and proliferative capacity (*25–28*). However, both micro-contact printing (25,26) and spheroid culture (27) restrict the proliferation of MSCs leading to limited or no expansion in cell number. On the other hand, creating nanopatterns (28) demands huge infrastructure and is cost.

In this work, we have shown that hMSCs maintain their stemness over long passages when cultured on an optimally soft poly-acrylamide (PAA) gel. Soft substrate also preserves cellular morphology. Staining for β-gal and BrdU showed respectively that in these cells onset of senescence is delayed and proliferative potential is maintained. Staining for other senescence related changes such as loss of Lamin B and gain of Lamin A confirmed this observation. Not only the proliferative potential but the cells cultured on gel could differentiate into adipo lineage, as shown by the expression of PPAR-gamma and accumulation of oil droplets while cells cultured on tissue culture plastic (TCP) lose their adipogenic differentiation potential. Finally, we have shown that surface markers, used to characterize MSCs, remain unaltered in the cells cultured on soft substrate ensuring the maintenance of cellular identity.

## Result

### Loss of cell morphology and induction of senescence during long-term in vitro expansion

To study the effect of substrate stiffness on maintenance of stemness, we cultured umbilical cord-derived hMSCs (UCMSCs) on polyacrylamide gel and on TCP, both coated with collagen I, from passage 3 (P3) to passage 13 (P13) (Fig. 1). These cells were well characterized (*SI appendix*, Fig. S1) and applicable bio-safety and ethical guidelines were followed. For better understanding of the long-term effect of passaging on cellular behavior, we grouped our results as *Early Passage* (EP), *Mid Passage* (MP), and *Late Passage* (LP) which were defined as passage number (P ≤ 6), (P = 7 to 10), and (P > 10), respectively. This classification, though arbitrary, was done based on the prevalent practice that MSCs are generally used till maximum P6 for research and clinical applications (10,29).

**Fig. 1:**
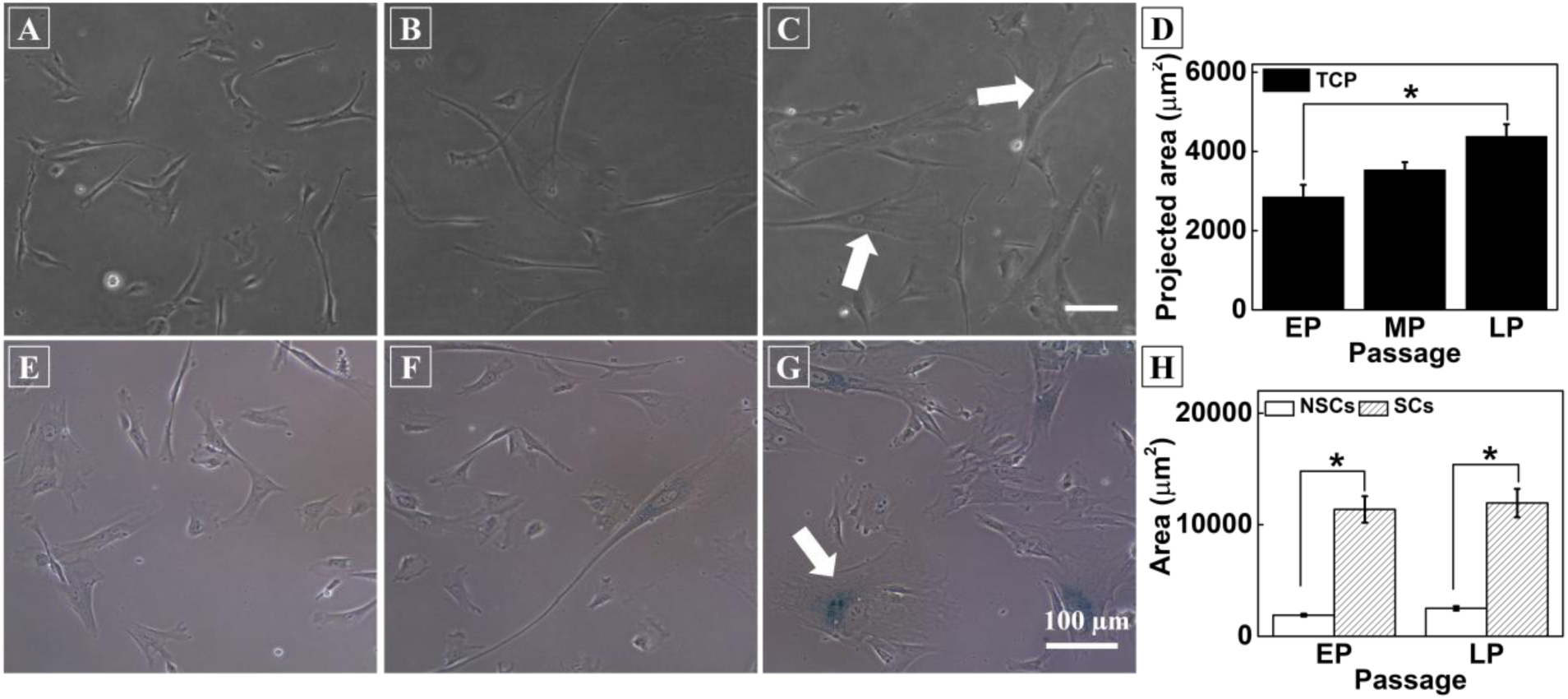
hMSCs lose their morphology and enter into replicative senescence on long-term passaging on tissue culture plastic (TCP). Representative micrographs from **(A)** Early passage (EP, passage number ≤ 6) **(B)** Mid passage (MP, passage number = 7 to 10) and **(C)** Late passage (LP, Passage number > 10) show that during *in vitro* expansion, MSCs lose their spindle morphology and become large and flat. White arrows show large cells with irregular shapes. **(D)** Over passage, average cell spread area increases significantly from ~3000 μm^2^ to ~4500 μm^2^. Data are shown from three independent technical replicates as mean ± SE (n = minimum 150 from each replicate). **(E-G)** β-gal staining (a senescence marker) significantly increased in MP and LP cells compared to EP cells. **(H)** Spreading area of senescent cells (SCs) and non-senescent cells (NSCs): Senescent cells have more spreading area than non-senescent cells, irrespective of passage (n = at least 100). Results are expressed as mean ± SE, statistics by t-test, *p<0.05.

To measure spread area, cells were imaged after 24 hours of cell seeding and average cell area was computed considering at least 150 randomly selected cells for each passage. We found that when cultured on TCP, UCMSCs with increasing passage lose their spindle morphology (Fig. 1 *A-C*) and go into replicative senescence (Fig. 1 *E-G*). Majority of the cells became flat and took irregular shapes with the passage as shown by the white arrows in Fig. 1 *C*. Also, more debris and more granularity in the cytoplasm were observed for later passage cells (data not shown). To check the onset of senescence, we trypsinized the cells from their respective substrates and re-plated them on glass coverslips. After 24 hours, we stained the cells with SA-β-gal, a well-established method to capture the senescent cells. We observed that while for EP cells, only very few cells (<5%) were β-gal positive (Fig. 1E), it increases gradually to finally reach at about 20% for late passage (LP) cells (Fig. 1 *G*). Further, to check if the increase in the cell area and increased senescence are associated, we estimated the average spread area of β-gal positive and β-gal negative cells for early as well as for late passage. We found that irrespective of passage number, senescent cells are always significantly larger compared to the non-senescent cells (Fig. 1 *H*). Also, the average area of senescent and non-senescent cells remains almost unaltered over passage implying a possibility of close association of cell spreading with the onset of senescence. This observation led us to the hypothesis that maintaining cell size using soft substrates made of polyacrylamide gel may delay or stop senescence resulting into more efficient *in vitro* expansion of MSCs.

### Selection of Substrate for further study based on cell morphology and proliferation

It is known from the literature that cells spread less and take round morphology on soft substrates (*30,31*). To find the suitable substrate that restricts cell spreading, we cultured LP cells on PAA gels of stiffness range varying from 0.5 kPa to 20 kPa and TCP (~GPa) (*SI Appendix*, Fig. S2). We found that on soft gels (E ≤ 5 kPa), LP cells had cell spread area smaller than or equivalent to EP cells cultured on TCP (Fig. 1D and *SI Appendix*, S2G). This observation implied that cell spread area can be kept restricted over passages if cultured on soft gels (E ≤ 5 kPa). However, the question remains how soft can we go? To answer this question, we need to consider another parameter i.e. the effect of substrate stiffness on cell proliferation. It is known that soft gel induces reversible cell cycle arrest or quiescence in hMSCs (24,32). As a result, the very soft gel cannot be used for cell number expansion. To find the optimum range of stiffness, we cultured cells on substrates of different stiffness. After 48 hours of culture on these gels, which is sufficient to induce quiescence (32), we gave a 4 hours pulse of BrdU which tags the replicating DNA. We found that while cells on 1 or 2 kPa gel showed critically less replicating DNA, cells on gels of 5 kPa and higher stiffness had more than 30% of dividing cells which is equivalent to that on TCP (*SI Appendix*, Fig. S2*L*).

Putting these two observations together, we selected 5 kPa gel for all our ongoing studies to compare the effect of substrate stiffness on long-term *in vitro* culture.

### Soft Substrate maintains Cellular morphology and Self-renewal ability of UCMSCs

When continuously cultured on 5 kPa gel for 33 days, it was observed that UCMSCs maintain their cellular morphology and proliferative potential better than the cells cultured on TCP. For P3 to P13, the cells were trypsinized in every 72 hours and re-seeded on the respective substrates at 1000 cells/cm^2^ seeding density. Cells cultured on gel are denoted with (/G) and the cells cultured on TCP are denoted by (/T). Our actin immunostaining data from different passage (Fig. *2A*) show that while cells lose their morphology when cultured on TCP for multiple passages, they maintain the same if cultured on the gel. However, to note that another substrate dependent phenotype i.e. focal adhesion shows an interesting trend. While there is a significant difference in the appearance of focal adhesion assembly for EP cells on gel and TCP, for LP cells they look identical (Fig. 2A). The average area of cells on 5 kPa gel was maintained with the passage at about 1500 μm^2^ but the same was increased from 3000 μm^2^ to 4500 μm^2^ on TCP (Fig. *2B*). On the gel, we observed fewer irregularities in cell shape as well. To have a blind test, we showed the cells under a microscope to multiple independent observers who were well versed with hMSCs morphology. They could not differentiate between EP/G and LP/G cells. However, the difference between EP/T and LP/T was obvious. Other than spread area, cells expanded on the gel for long passage also showed significantly lower traction force (Fig. 2C) than the cells on TCP when tested on 5 kPa gel for traction force microscopy (TFM).

**Fig. 2.**
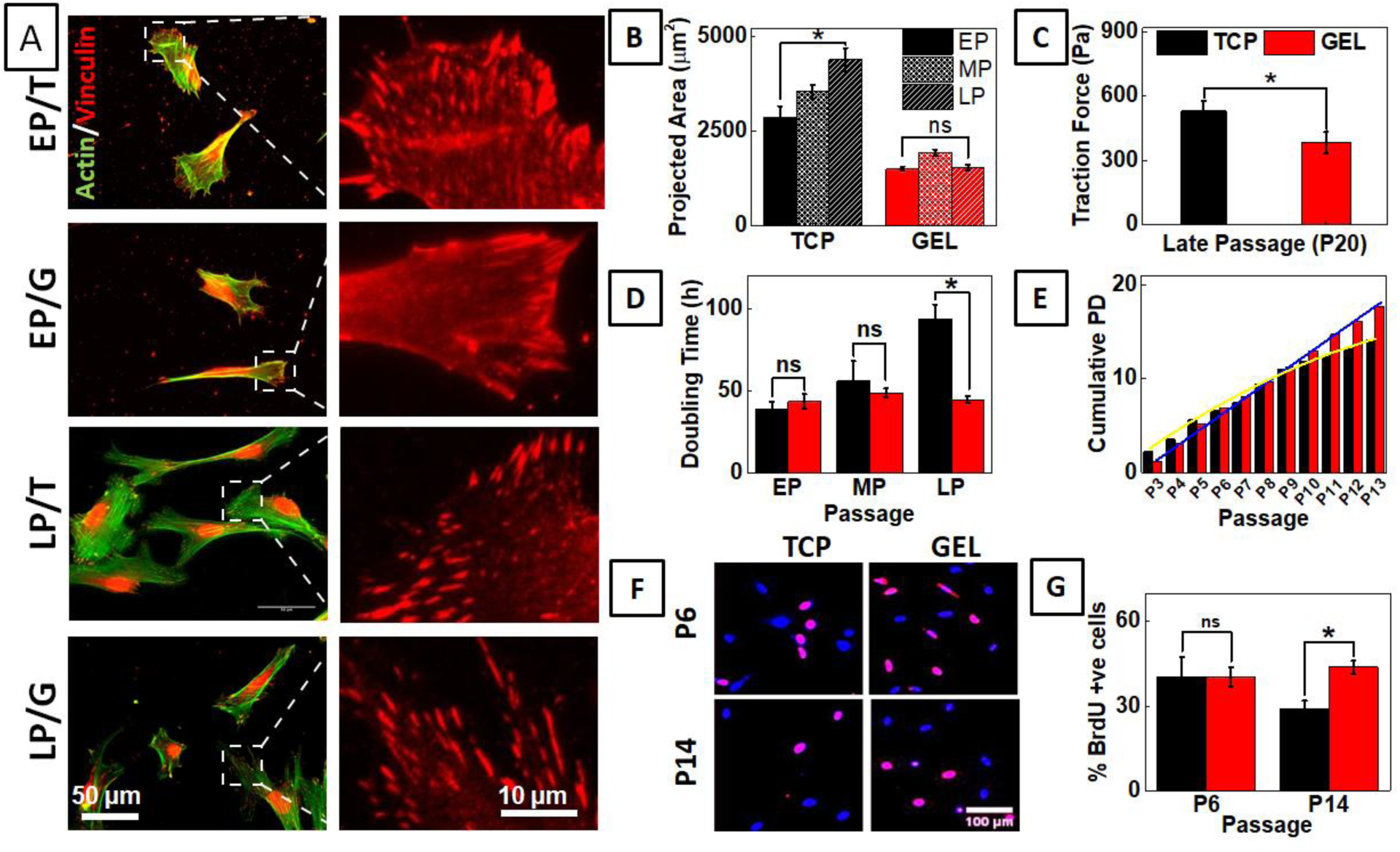
Soft substrate maintains cellular morphology, rate of expansion and proliferation during serial passage. (A) Early (EP) and late passage (LP) UCMSCs were serially cultured on soft gel (G) with collagen-coated TCP (T) as positive control. Morphological changes during long-term expansion was examined by immunostaining of actin (green) and vinculin (red). Late passage cells cultured on gel showed maintained morphology. (B) Cell spread area increases with passage on TCP; however spreading area of cells on gel showed no significant difference across passages. (n = 3, N = 150, *p < 0.05) (C) Late passage UCMSCs on gel showed significantly lower traction than that on TCP. (D) Doubling time of UCMSCs cultured on gel and TCP over the passage. While doubling time increases with passage when cultured on TCP, the same remains unaffected when expanded on 5 kPa gel. The difference in doubling time for EP and MP are negligible while for LP the difference is significant. (E) Cumulative population doubling (CPD) of UCMSCs cultured on gel and TCP over passage. CPD increases linearly for UCMSCs on the gel but approaches a plateau for cells cultured on TCP. (F) Immunofluorescence images of nuclei co-stained with DAPI (Blue) and BrdU (Red) capturing relative percentage of cycling cells. (G) Percentage BrdU positive cell for P14 (LP) from gels are significantly higher than the cells from TCP though the BrdU uptake was same for P6 (EP) cells. This data reconfirms that cells maintain their proliferative potential if cultured on the soft gel. Results are expressed as mean ± SE. *p < 0.05, scale bar = 200μm, n = at least 150 from each passage, total at least 450 for each condition.

To check if along with maintenance of morphology, self-renewal efficiency is also maintained, we assessed the effect of soft substrate on cellular expansion of UCMSCs in the long-term passage. Using microscopic images as described in the method section, we counted cell number twice; once, after 4 hours of seeding and second, just before harvesting. From the number of cells seeded and harvested, we calculated population doubling (PD), doubling time (DT) and Cumulative PD (CPD) as described in methods section. Fig. 2*D* compares DT values for EP, MP and LP cells. We found that doubling time (DT) for cells on TCP increased slightly from 40 h to 50 h for EP to MP and then jumped significantly to more than 80 h for LP cells. However, DT value remains almost constant for all three groups of cells (~ 40 h) when cultured on 5 kPa PAA gel.

This observation is also reflected in cumulative population doubling (CPD) (Fig. 2E). We observed that CPD on TCP increased almost linearly till P9 and then gradually slowed down. For gel, CPD continues to increase linearly till the end of the experiment. By P20, we observed a difference of 9 in CPD signifying 2^9^ or 512 times more UCMSCs can be obtained from gel than from TCP. This difference signifies that while one single cell will give us 4 million cells after 20 passages if cultured on TCP, the same cell will give 2000 million or 2 billion cells if cultured on the soft gel.

Further, to confirm that this reduction in CPD on TCP compared to that on the gel is indeed due to the maintenance of proliferation and not a cell culture artifact, we compared the fraction of cycling cells present in the cells harvested from gels as well as from TCP after reseeding them on the glass. As expected, we found that while there was no significant difference between EP/T and EP/G cells, the percentage of S phase cells in (LP/G) was significantly higher than that in (LP/T) cells (Fig. 2*F*-G). This data conclusively proves that 5 kPa PAA gel maintains self-renewal ability of UCMSCs over long-term culture leading to significant increase in cell number. To rule out the possibility of higher cell adhesion on gel compared to TCP resulting into higher CPD, we checked for plating efficiency on both the substrates and found no significant difference (*SI Appendix*, Fig. S3).

All the result mentioned here were done with technical triplicates. To check the reproducibility of the data, we used two independent biological replicates and MSCs derived from bone marrow. We found the similar observations in all of these cases, as shown in *SI Appendix*, Fig. S4-S8.

### Soft substrate delays senescence

To verify, if this maintained rate of expansion on soft gels is a result of reduced senescence as we proposed earlier, we stained the cells with SA-β-gal. We found that on TCP, fraction of senescent cells increases with the passage, reaching >20% of the population for late passages (Fig. *3A-C* and *G*). However, when cultured on the gel, although there was an increase in senescent population from early to mid-passage, the percentage of this population did not increase further and remained <10% which is significantly less compared to the same for cells cultured on TCP (Fig. *3D-F* & *G*). To reconfirm, we also checked for expression of vimentin, which is known to over express in senescence fibroblast (33,34). Immunofluorescence analysis revealed a staggering 4 fold increase in vimentin expression in cells cultured on TCP compared to the same on gel (*SI Appendix*, Fig. S9).

**Fig. 3.**
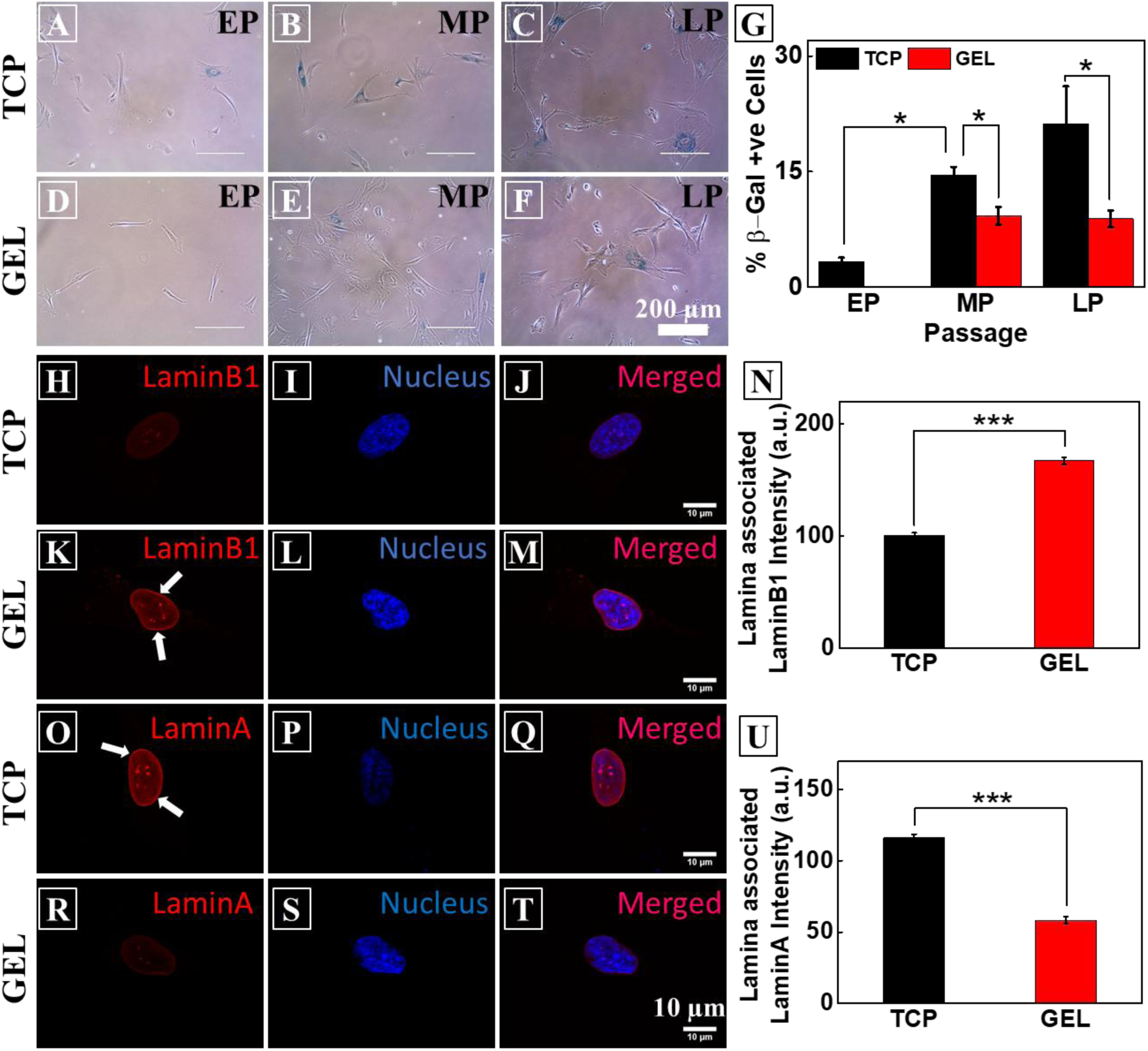
Soft substrate delays senescence. (A-G) Cells, stained for β-gal (blue), a well-established senescence marker, show that while senescence gradually increases for TCP over the passage, on the gel after an initial increase, it remains constant. The same is quantified in figure (G). Minimum 200 cells per passage were analyzed. (H)-(N) Comparison of expression of LaminB1(red), at the nuclear periphery/lamina. The decrease of Lamin B1 expression on TCP (H-J) compares to UCMSCs on the gel (K-M). Quantification of intensity shows that Lamin B1 is maintained on late passage UCMSCs on the gel (N). Further, comparison of Lamin A expression for late passage UCMSCs on plastic(O-Q) and on the gel (R-T) reveals the fact that Lamin A expressed more on TCP. The same is quantified in figure (U). Minimum 10 cells were used for analysis with at least 5 positions on the periphery of each nucleus in figure (N) and figure (U). Results are expressed as mean ± SE *p<0.05, ***p<0.001

We also checked for Lamin A and Lamin B1, two nuclear envelop associated proteins, expression of which is known to vary differentially in senescence (22,23). We trypsinized cells from gel and TCP and plated them on glass coverslips. After 24 hours of cell seeding, we stained the cells with respective antibody and quantified Lamin A and B1 expression on the lamina of the nucleus. We indeed found that while expression of Lamin A is more in the cells cultured on TCP (Fig. *3O-U*), these cells lose the expression of Lamin B1 (Fig. *3H-N*). Both of these observations, along with SA-β-gal staining data, show that culturing UCMSCs on soft gel delays senescence.

### Long-term culture on gel did not alter surface marker expression but helps the stem cells to maintain differentiation potential

Finally, to confirm the identity of our cells after long-term culture, we investigated the effect of substrate on the expression of surface pluripotency markers. Surface marker analysis of late passage (P22) UCMSCs using flow cytometry and immunocytometry demonstrated that cells cultured either on gel or on TCP express characteristic positive surface markers, CD105 (Fig. 4*A* and *E*), CD44 (Fig. 4*B* and *F*), CD90 (Fig. 4*C* and *G*) and Stro-1 (Fig. *4D* and *H*). This is in accordance with the previous studies which showed that the MSC surface marker expression remains same with increasing passage (35). However, the difference appears in their differentiation potential. While LP/T cells lose their adipogenic potential, LP/G cells did not. After culturing both types of cells on TCP and in adipogenic induction media for 7 days, the cells were fixed and stained for the early adipo transcription factor, PPAR-γ. It is evident from the images that LP/G cells expressed a much higher level of PPAR-γ than the LP/T cells in similar condition (Fig. 4*I-K*). The same difference was observed after 14 days of adipo induction in terms of accumulation of oil droplets that was detected with Oil red O (ORO) dye (Fig. 4*L* and *M*). Approximately 75% of LP/G cells differentiated into adipocytes as determined by ORO in comparison to only 18% of LP/T cells staining (Fig. 4*N*).

**Figure 4:**
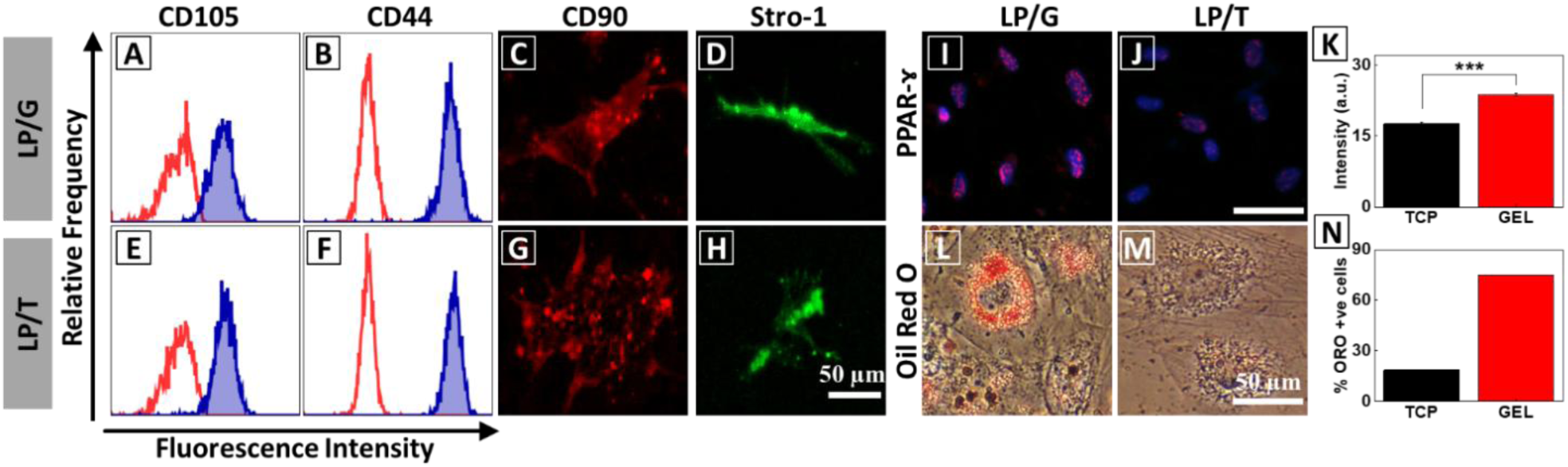
Long-term culture on gel did not alter surface marker expression but helps stem cells maintain differentiation potential. Flow cytometry analysis of the expression of surface pluripotency markers of UCMSCs at the late passage (P22) from the gel (LP GEL) (A-D) and TCP (LP TCP) (E-H) was determined. The expression of surface markers, CD105, CD44, CD90, and Stro-1 was not altered by the long-term culture of UCMSCs on the gel. Late passage hMSCs from gel and TCP were cultured in adipogenic induction media for 14 days. **Differentiation markers**: The early adipogenic marker, PPAR-γ, expression was checked using immunostaining of hMSCs from LP GEL (I) and LP TCP (J) (magenta puncta in blue nucleus). The UCMSCs serially passaged on gel showed higher expression of PPAR-γ compared to the cells cultured on TCP as quantified in figure (K). (L-O) After 14 days in adipogenic induction media, the LP gel UCMSCs accumulated the lipid droplets, stained with oil red O (ORO) (L) whereas the lipid droplets were not accumulated in LP TCP hMSCs (M). The % of ORO positive cells were more on gel substrate compared to TCP as quantified in figure (N). PPAR-γ intensity was quantified using 273 nuclei and plotted as mean ± SE. ***p < 0.001

## Discussion

In this work, we have demonstrated that a soft PAA gel substrate of 5 kPa elastic modulus can maintain the stemness of UCMSCs *in vitro* by delaying the senescence process. Like any other primary cells, hMSCs undergo only a certain number of divisions before entering into replicative senescence. Replicative senescence is typically marked by large and flat morphology, and significantly slowed down proliferation rate (8,9). Multi-lineage differentiation potential also goes down with population expansion (10,11). This decreased proliferation and loss of multi-lineage potential poses a serious challenge in using *in vitro* expanded hMSCs for clinical use. In this work we have shown that culturing UCMSCs on soft substrate maintains self-renewal ability, multi-lineage potential and surface markers much beyond the same when cultured on TCP.

As previous studies have shown that span of culture before onset of senescence program can vary significantly depending on cellular origin, we first cultured our UCMSCs on TCP up to 13 passages and confirmed the onset of senescence with enlarged and flat morphology, and β-gal staining (Fig. 1) (36,37). We also showed that average area of non-senescent cells remains same over passages. This observation opens up a possibility that restricting cell spreading may help in delaying senescence. Such hypothesis is supported by the work of Killian et al. which showed that MSCs cultured on micro-patterned substrates of restricted spread area helped in maintaining the expressions of stemness markers STRO-1 and Endoglin (26). Although stemness was maintained, such approach is not very useful for MSC expansion as cells on micro-island stop proliferation and do not increase in number.

Earlier studies also showed that stemness in hMSCs could be maintained better by reducing acto-myosin contractility using pharmacological inhibitor of ROCK and myosin (25). In a different study, it was shown that while mESCs (mouse embryonic stem cells) grown on TCP needs LIF to stop spontaneous differentiation, a very soft substrate (600 Pa) can maintain them in their undifferentiated state without LIF (38). All these results indicated that cell area/contractility plays an important role in loss of stemness and both can be kept low if cultured on a soft substrate (31,39). However, none of these works checked the effect of substrate stiffness in long-term culture. To the best of our knowledge, this is the first work to demonstrate the effect of substrate stiffness on long-term culture for any cell type.

Consistent with previous report, we also found that though UCMSCs lose their self-renewal ability when cultured on TCP, they maintain the molecular signatures related to stemness (Fig. 4A-H)(40). The cells, irrespective of cultured on gel or TCP were positive for CD105, CD44, CD90, and Stro-1. However, cells cultured on TCP lost their adipogenic differentiation ability whereas the same was maintained for the cells cultured on gel (figure 4I-N). Loss of adipogenic potential over long term passages and dominance of osteogenic differentiation has been reported by many earlier researchers (41,42,). It was shown that flat cells that appear spontaneously over long term culture, lose their adipogenic potential (40). This observation is not unexpected if we look from cell mechanics angle. It was established by Engler et. al. in their seminal paper in 2006 that stiff substrate (34 kPa) induces osteogenic lineage in hMSCs (21) even in absence of chemical inducer. Similarly, it was also shown that cells that were made to spread more or to take the shape that induces high contractility, also were prone to osteogenic lineage commitment (39,43). So, it is expected that if for multiple passages, cells are continuously exposed to a substrate as rigid as TCP which increases cellular spreading and contractility, adipogenic potential would get diminished. However, a soft culture substrate, in contrast, should maintain the multi-lineage potential, as demonstrated by our result (Fig. 4).

Other than UCMSCs, we also used bone marrow derived hMSCs and found the similar result proving that this effect might not be source specific. However, we also found that the substrate stiffness for optimal growth of skin derived keratinocytes is not the same as for MSCs (data not shown). How cell type and optimum substrate stiffness are inter-linked is open for future investigation.

One of the interesting observation in this work is that soft substrate delays senescence. It is known that acquiring replicative senescence over in vitro expansion may not be an obvious purposeful program but a result of the external environmental condition. For example, it has been shown that increased oxygen may induce senescence faster. In contrary, the hypoxic condition is known to maintain stemness for hMSCs (44). However, the effect of substrate stiffness on senescence has not been studied before. We have demonstrated using four known markers of senescence namely expression of β-gal, loss of Lamin B1, gain of Lamin A and vimentin (10,45,46) that an optimally soft substrate may delay the onset of senescence significantly.

In summary, our data show that instead of using TCP, culturing cells on the soft substrate will help to solve the problem of limited availability of MSCs, by increasing the number of available cells after extended expansion. This work offers a possibility to design cell-specific culture substrate in future. This work also demonstrates for the first time that replicative senescence in hMSCs can be delayed using substrates of physiological stiffness.

## Experimental Section: Methods

Bone marrow derived human MSCs were purchased from Lonza, and fresh umbilical cord were obtained from healthy individuals after due ethical clearance and bio-saftey approval. Detailed information on methods, cell culture, immuno-staining, are presented in *SI Appendix*.

## Supplementary Materials

**Figure S1:**
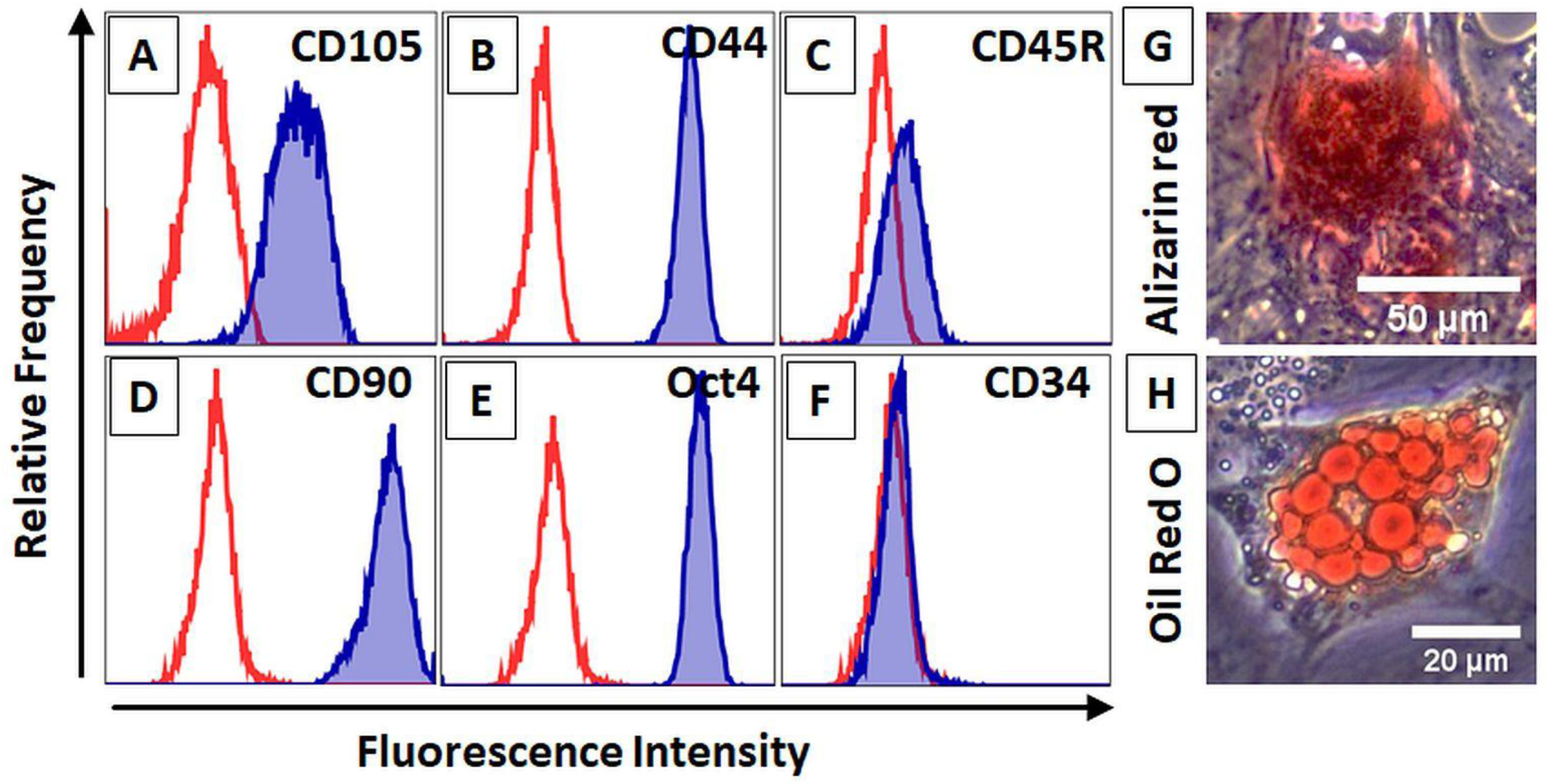
Characterization of hMSCs: Flow cytometry analysis of the expression of positive and negative markers of UC-hMSCs at early passage (P4) was determined (A-F). The expression of surface markers, CD105 (A), CD44 (B), CD90 (D) and pluripotency marker, Oct4 (E) was positive whereas the cells did not express CD45R (C) and CD34 (F) which are negative markers for hMSCs. Red line in flow cytometry data is the auto-fluorescence and the blue filled histogram is the fluorescence signal. hMSCs were cultured in osteogenic and adipogenic induction media for 21 days and 14 days, respectively. Alizarin red staining was performed to identify the calcified nodules in the osteogenic hMSC (G). The adipogenic hMSC accumulated the lipid droplets stained with oil red O (H).

**Figure S2:**
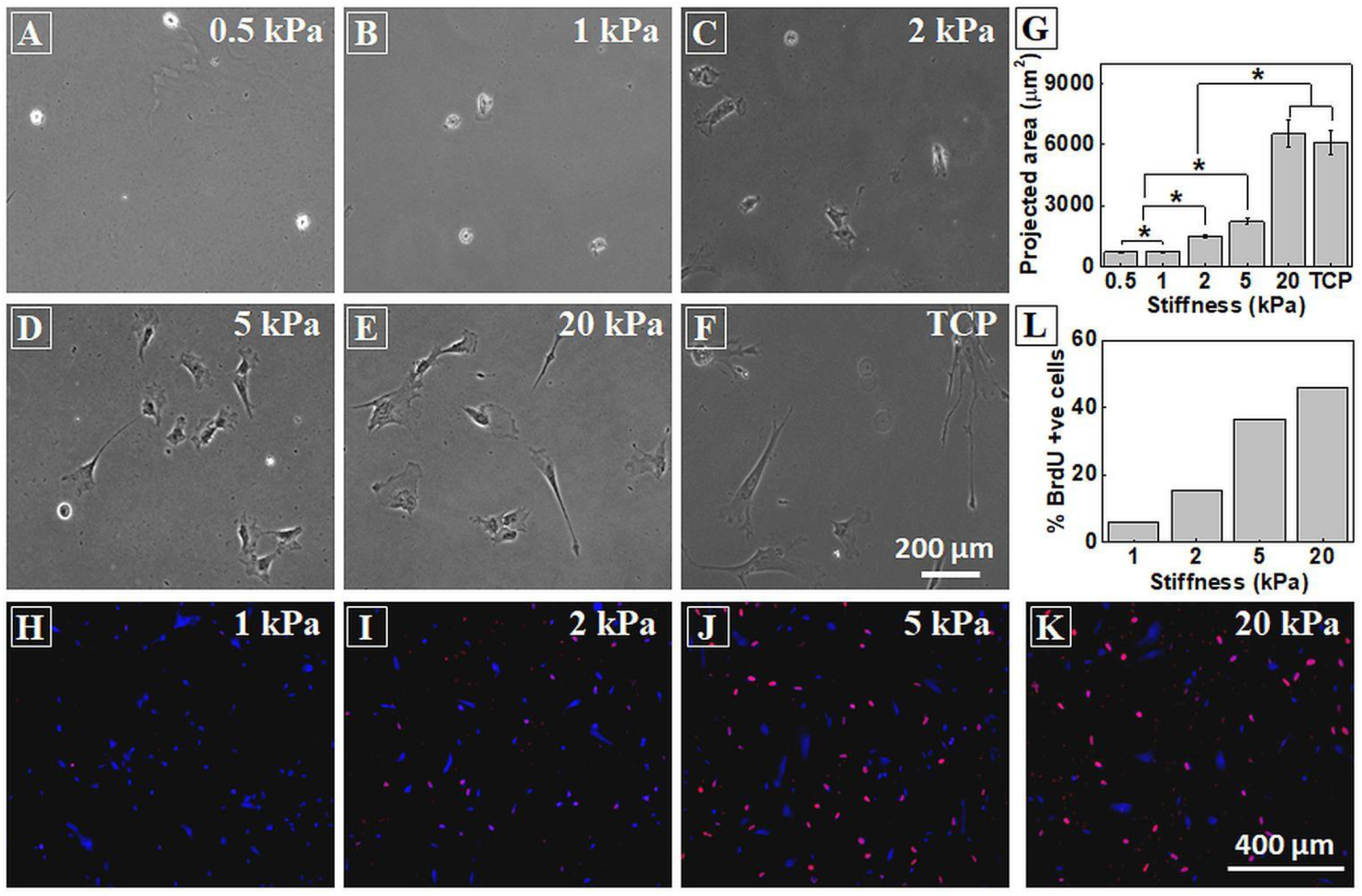
Soft Substrates restrict cell spreading and hinders proliferation: Phase contrast images of LP UCMSCs after 24 hours of seeding on substrates of various stiffness, (A) 0.5 kPa, (B) 1kPa, (C) 2 kPa, (D) 5kPa, (E) 20 kPa, and (F) plastic (TCP) show that cell spread area increases with substrate stiffness. (G) Quantification of average spreading of LP cells on various substrates shows that LP cells on 5 kPa and lower are even smaller than the EP cells on TCP. Average area of EP cells on TCP is shown by the dash line. Scale bar=200μm, 150 cells were taken for area analyses, results are expressed as mean and s.e.m, statistics by unpaired t-test, **p<0.05.(H-K) Representative images of BrdU positive nuclei (pink) on PAA gel of various stiffness (1,2,5, and 20 kPa) and (E) their quantification show that very soft gel prohibits cell proliferation. Min 250 nuclei were counted from random 15 images for analysis. Pink: BrdU positive nuclei, Blue: DAPI, BrdU negative nuclei. Graph suggest that hMSCs are more proliferative on gel of 2 kPa and above. Data are expressed in mean ± SE.

**Figure S3:**
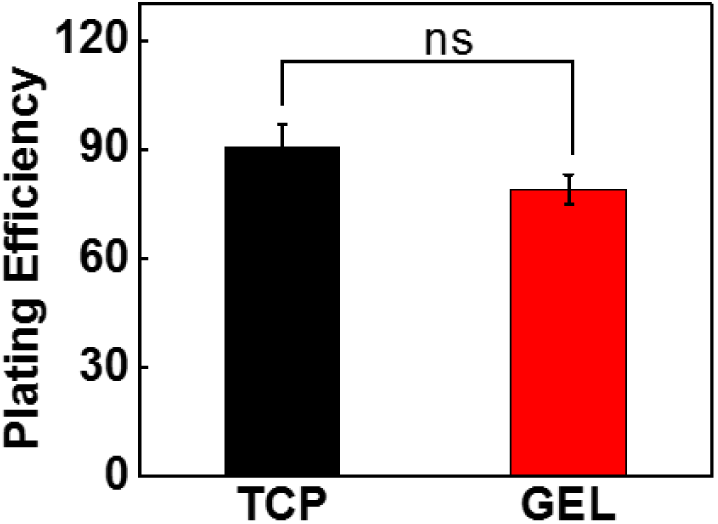
Comparison of Cell plating efficiency between 5kPa gel and TCP. Plating efficiency determined by no. of cells seeded/ no. of cells attached *100. Results are expressed as mean and s.e.m, statistics by unpaired t-test, **p<0.05. with three technical replicates.

**Figure S4:**
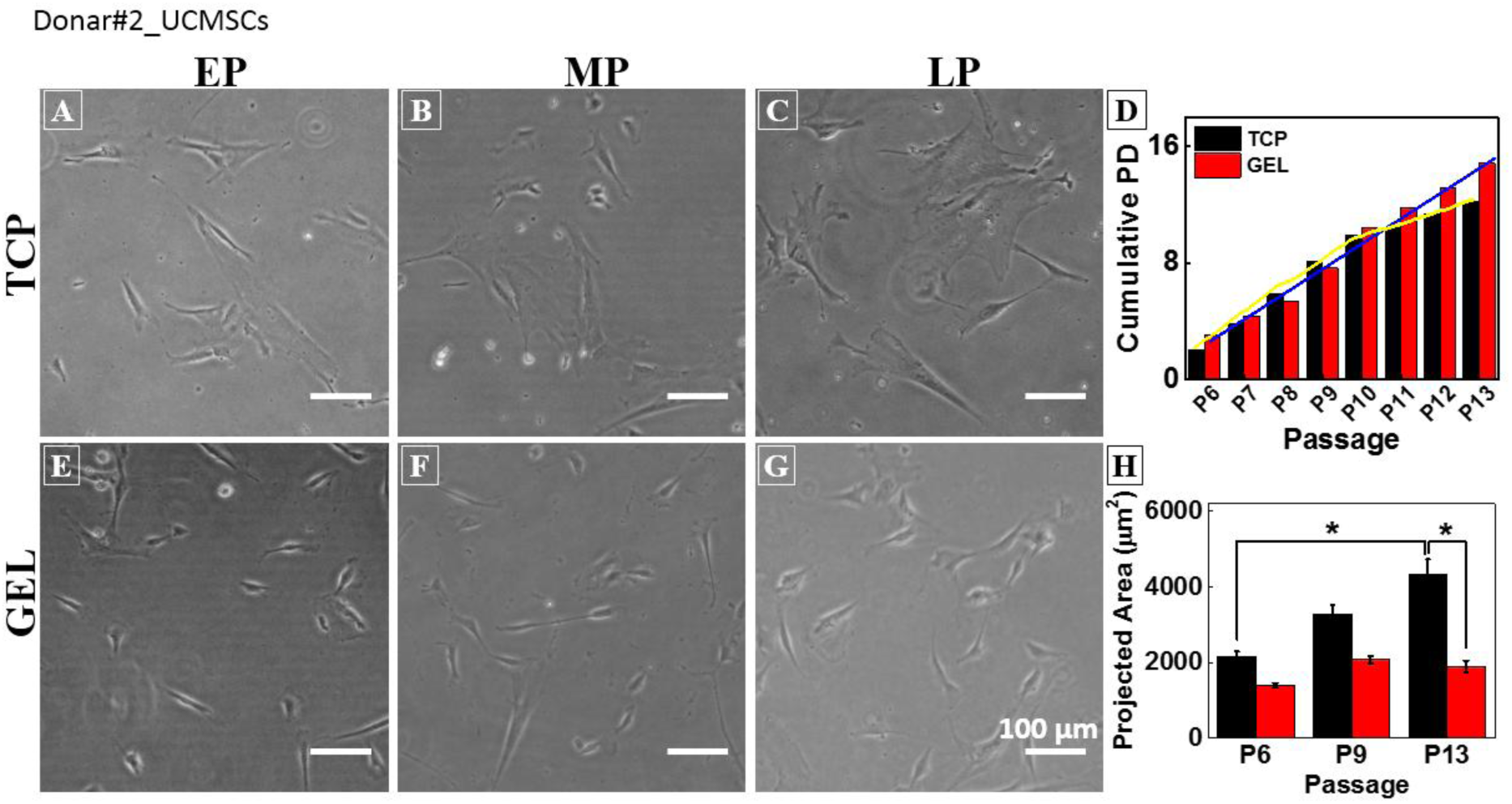
Biological repeats of data using UCMSCs of donor#02. (A-C) Representative phase contrast images of UCMSCs of donor#02 on TCP and (E-G) on GEL. (H)Spread area increases from 2200 at EP to 4400 μm2 at LP. Whereas spreading remain 2000–2300 μm2. (D) Cumulative PD plot against passage number. Results are plotted mean±s.e.m. statistics by unpaired t-test, *p<0.002, n=3, and n=150.

**Figure S5:**
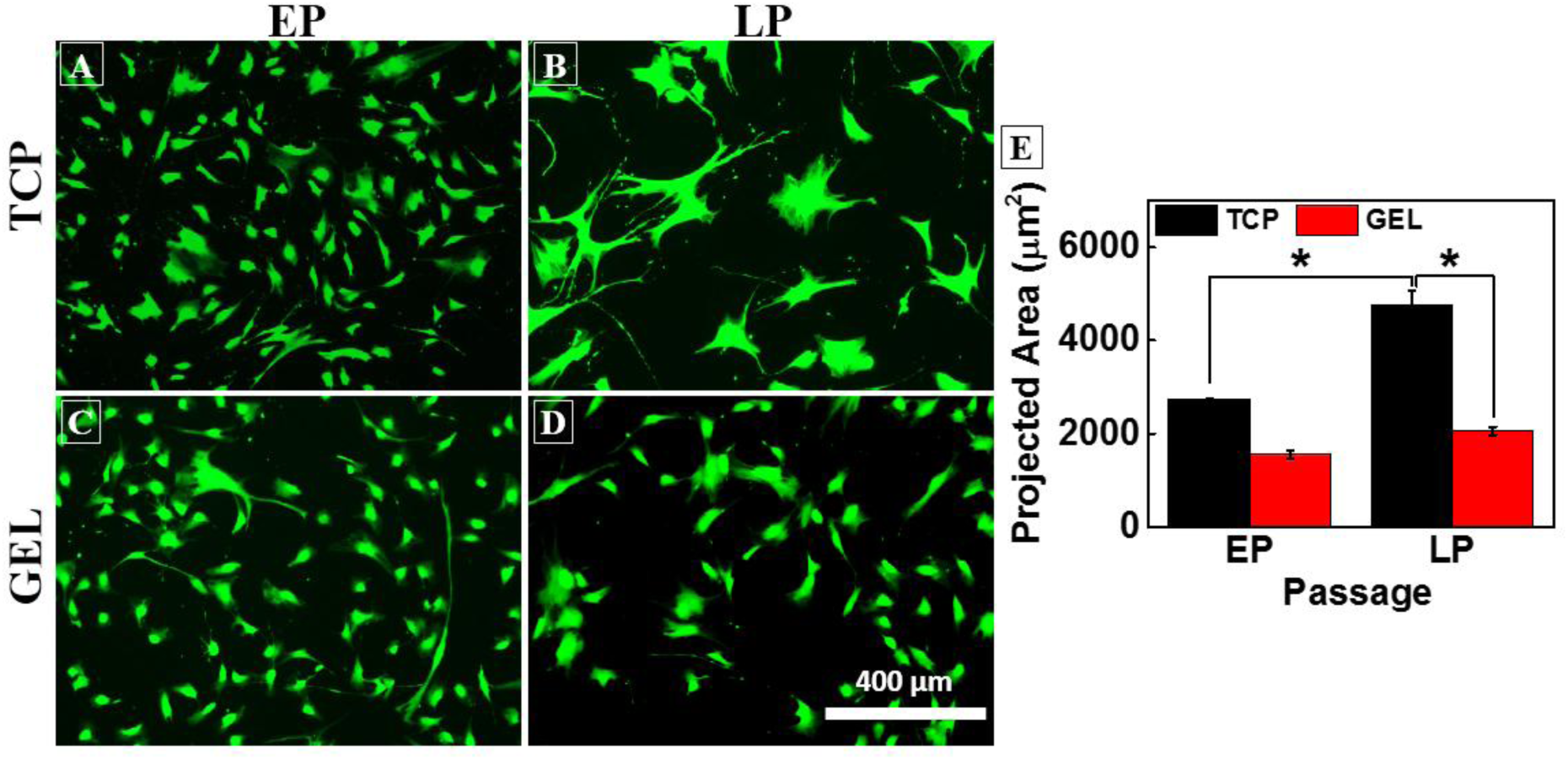
Morphological changes of bone marrow derived hMSCs. (A-B) immunofluorescence imaged of cells taken from TCP, (C-D) from GEL. (E)Quantification of area data shows that Soft gel restrict spreading while cell spread on TCP from 2500 to 5000 μ*m*^2^.results are expressed in mean±s.e.m.** p value <0.05 using student t-test, (n=3 N=150).

**Figure S6:**
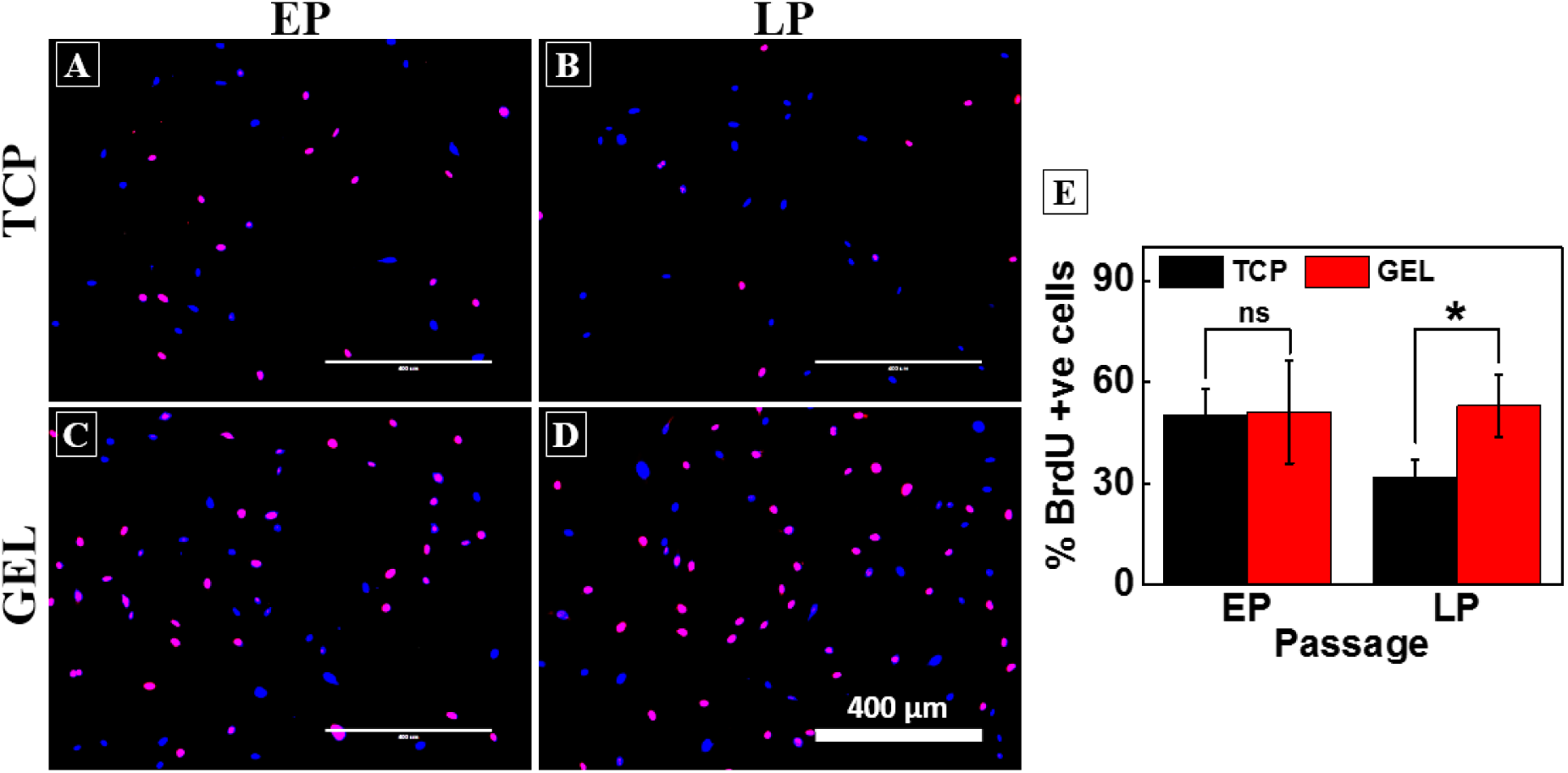
DNA replication of bone marrow derived hMSCs. BrdU study shows Compared to TCP(A-B) soft gel maintain DNA synthesis(C-D). Blue dots show nucleus and pink dots represent cells in synthesis phase. (E) Percentage of pink coloured cells (S-phase cells) are determined by counting cells manually. Around 400 nuclei with three experimental replicates. results are expressed as mean and s.e.m, statistics by unpaired t-test, **p<0.05

**Figure S7:**
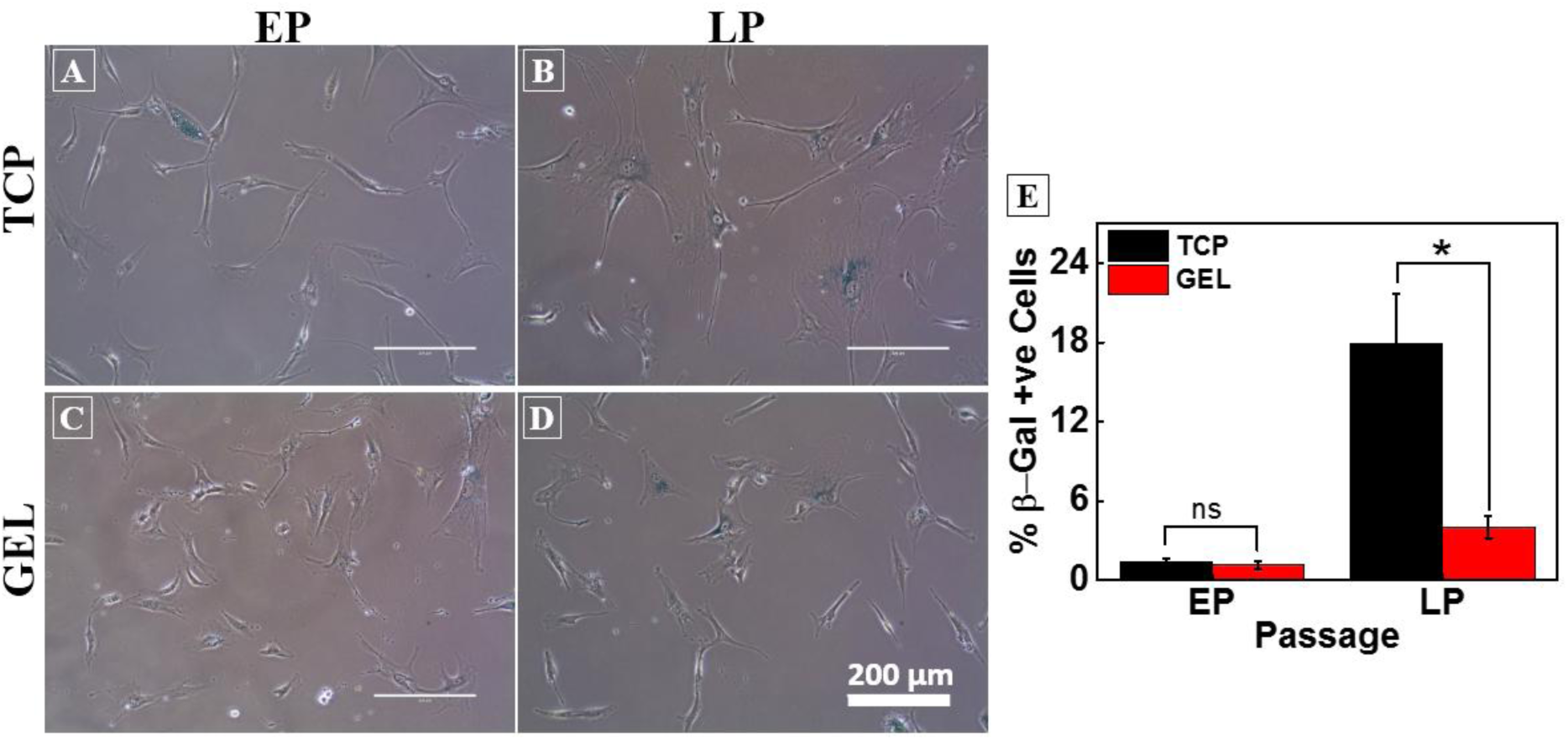
Replicative senescence of bone marrow derived hMSCs. Late passage cells become senescent cells. Beta gal associative assay is used to identify senescent cells. blue coloured cells are senescence cells. Coloured images show that soft gel (C-D) have less number of beta gal positive cells than (A-B) TCP in late passage. (E) Min 150 cells were counted manually from random 20 images for quantification. Results are expressed as mean and s.e.m, statistics by unpaired t-test, **p<0.05.

**Figure S8:**
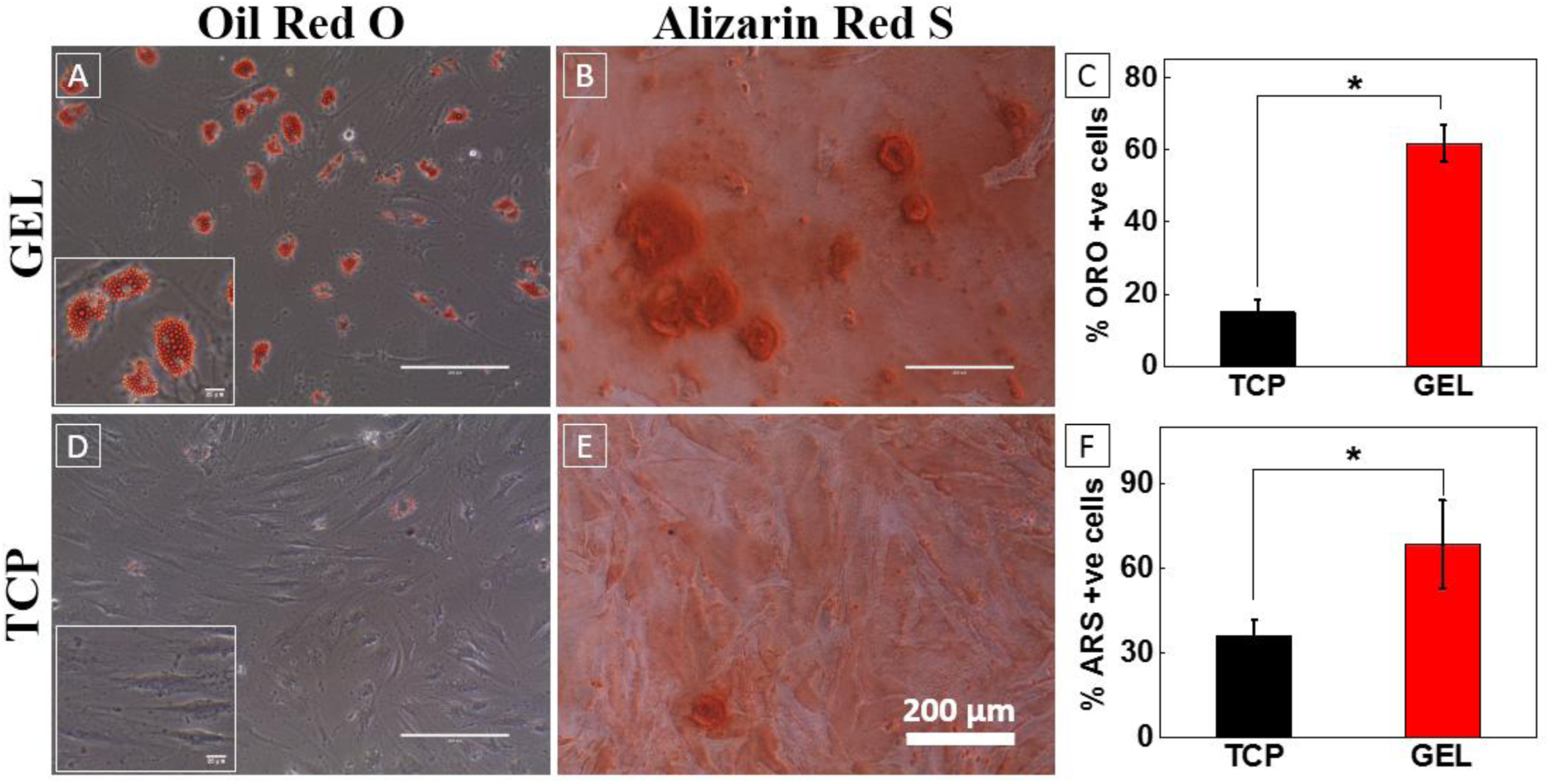
Oil red O stained coloured images (A &D) and Alizarin Red S stained images (B&E) for accumulation of lipid droplets and calcium deposition of late passage bone marrow hMSCs. (C) Quantification of oil red o was done base on number of adipocytes by counting cells manually. Similarly, calcium deposition was determined using manually osteoblast cells. Results are expressed as mean and s.e.m, statistics by unpaired t-test, **p<0.05. Minimum 100 cells were counted from 10 random images.

**Figure S9:**
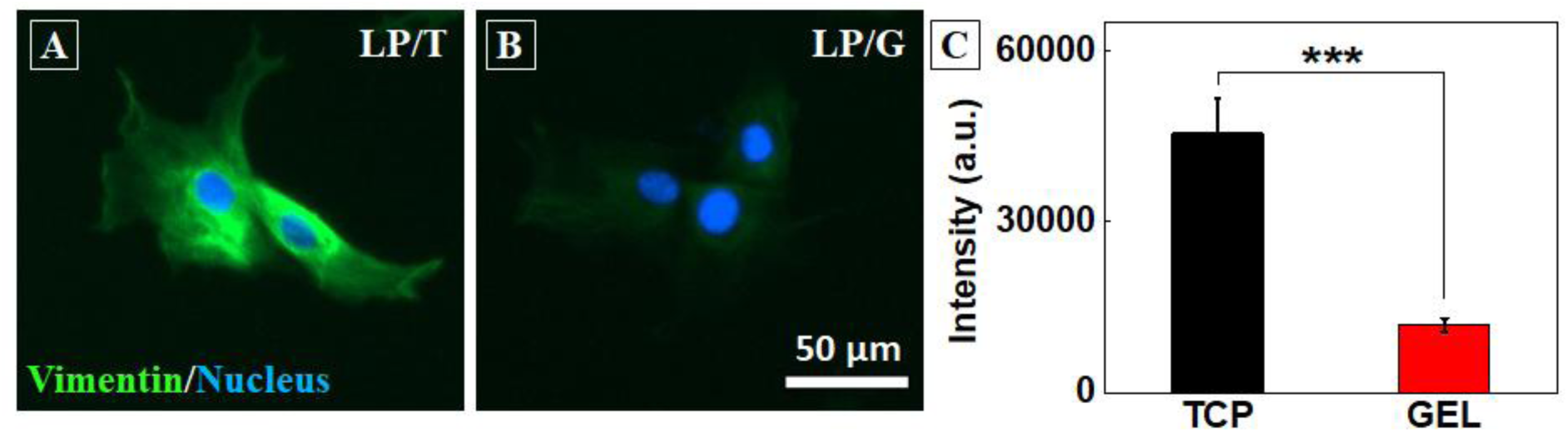
High expression of Vimentin in LP/T cells. (A and B) UCMSCs from TCP (A) and gel (B) were stained for Vimentin, a known marker for senescence. (C) LP/T cells express significantly higher vimentin compared to LP/G cells. Results are expressed as mean ± SE ***p<0.001

## Materials and Method

### Substrate preparation

Gels of polyacrylamide (PAA) of various stiffness were prepared by mixing 40% polyacrylamide and 2% bis-acrylamide solution, as described previously (*1*). Substrate preparation protocols and modulus values were adopted from previously published work (2). Briefly, the gels solution for desired stiffness was mixed with APS (ammonium persulfate) 1:100 and TEMED (1:1000) and placed between a hydrophobic glass (octadecyl-trichlorosilane (Sigma-Aldrich;**104817**) treated) and the transparency sheet 3-APTMS (Alfa Aesar;A17714) treated. Once polymerized, the hydrophobic plate was carefully removed. The gels were conjugated with sulfo-SANPAH and incubated with rat tail type I collagen (25 μg/ml) (Invitrogen;A1048301) at 4°C for overnight, as described (3). The tissue culture plates (TCP) (control) were also coated with type 1 collagen (25 μg/ml). The thickness of the gels was controlled by using defined volume of gel solution throughout the experiments.

### Cell culture

For serial passage experiments, P4 cells were seeded on large area gels and on TCP (both collagen-I coated as mentioned above) with same seeding density (1000 cells/cm^2^) in MSCs qualified medium α-(MEM)(Invitrogen:A1049001)/Low-glucose DMEM supplimented with 16% MSC certified Fetal Bovine Serum (FBS) (Invitrogen;12662029), 1% Glutamax (Invitrogen;35050061), and 1% pen-strep (Invitrogen;15140122) in humidified incubator with 37°C and 5% CO_2_. After 72 hrs of culture, cells were trypsinized from PAA gels and TCP using TrypLE™ Enzyme Express (Invitro;12604013) and were reseeded on fresh respective substrates and cultured for next passage, this process was repeated until the TCP growth halted.

### Cell count using image analysis

Images of the gels and TCP were acquired using Magnus™ microscope after 4 and 72 hr of seeding to determine accurate cell number for calculating population doubling (PD) as described in (4). For PD counting, 20 random images per sample were captured (covering ~3% of total area of gel), the average number of cells per frame was obtained and then divided by the total area of the frame to obtain seeding density (cells/cm^2^). The seeding density was then multiplied by the total area of the substrate (gel 20 cm^2^; TCP 25 cm^2^) to get the total number of cells seeded (4 hr) and harvested (72 hr) from a particular experimental condition (PAA gels and TCP) for respective passage. This was done at every passage, which was then used to calculate cumulative population doubling (CPD) for each experimental condition. Quantification of Cell morphology:

Cell images were captured at different passages at 48 hr post seeding using EVOS-FL auto inverted microscope (Life technologies, USA) at 10X magnification. Cell spreading area was determined using ImageJ (National Institutes of Health, Bethesda, USA) software by manually tracing around the perimeter of an individual cell. For each sample minimum 150 random cells were analyzed.

### BrdU assay

To check the percentage of S-phase cells in the cell cycle, cells from passage early passage (EP), mid passage (MP), and late passage (LP) were trypsinized from gels and TCP, and were seeded on collagen-coated glass coverslips as described above. After 48 hr of seeding, BrdU reagent (Invitrogen;000103) was added in 1:100 (v/v) ratio in media and incubated for 4 hr at 37°C humidified incubator with 5% CO_2_. Thereafter cells were fixed, permeabilized, blocked, and incubated in anti-BrdU antibody (Invitrogen;B35128) and counter stained with AlexaFluor 568 (A11061). Immuno-fluorescence images were captured using EVOS-FL auto and BrdU positive and negative cells were counted manually using ImageJ.

### Senescence assays

Senescence-associated β-galactosidase (SA-β-gal) was used to detect MSCs senescence using SA-β-gal staining kit (Abcam;AB65351) according to manufactures instructions. Briefly, cells from EP, MP, and LP were seeded in 6-well plate and incubated in growth media for 48 hr. Afterward, cells were fixed, stained with β-gal solution and incubated at 37°C without CO_2_. 10–15 random images were captured for each condition for analysis β-gal positive cells were counted manually.

### Differentiation assays

EP and LP cells from gel and TCP were seeded in a 12-well culture plate in growth medium for 72 hr and then incubated with differentiation media for adipogenic and osteogenic differentiation (Invitrogen;A1006501; A1006601 respectively) as per manufacturer’s manual. MSCs cultured in growth media were used as negative control. Post 14 days and 21 days incubation for adipo and osteo differentiation respectively, osteoblast were assessed with Alizarin Red solution (Sigma Aldrich;3422613022311) and adipocytes were assessed with Oil Red O (Sigma Aldrich;O0625) solution prepared as per manufacturer’s manual. Images were captured for qualitative and quantitative analysis using EVOS FL Auto.

### Immunofluorescence staining

For nuclear lamins (Lamin A, LaminB1), focal adhesion (FAs), and early adipogenic differentiation marker staining, EP and LP cells from gel and TCP were cultured on collagen-I coated glass coverslips for 24 hr. Cells were then fixed with 4% paraformaldehyde (PFA) in PBS for 15 min at room temperature (RT) and blocked (3% Bovine Serum Albumin in PBS) for 30 min and washed with cytoskeletal buffer (CB), as described previously (3). Cells were incubated with respective primary antibody (1:500, if not otherwise mentioned) for 4 hr at 4^0^C, and then incubated with corresponding secondary antibody for 1 hr at RT. Primary and secondary antibodies were used in following combinations: Alexa Fluor Phalloidin 488 (Invitrogen; A12379) (1:50 in blocking solution), anti-vinculin (Sigma Aldrich;V9131) counter stained with AlexaFluor 568 (Invitrogen, A11061), anti-PPAR-γ (Abcam; ab59256) counter stained with AlexaFluor 488 (Invitrogen; A11034), anti-LaminA (Abcam; ab8980) counter stained with AlexaFluor 568 (Abcam; ab175473), anti-LaminB1(Abcam; ab16048) counter stained with AlexaFluor 568 (Abcam; ab175470), anti-Vimentin (Sigma Aldrisch; V5255) counter stained with AlexaFluor 488 (Invitrogen; A11059). All secondary antibodies were used in 1:1000 dilution in CB. Cell nuclei were stained with Hoechst 33342 (Invitrogen;H3570) (1:1000) in PBS for 5 min at RT and mounted. Images were captured for qualitative and quantitative analysis using EVOS fluorescence microscope (Invitrogen USA).

### Traction force microscopy (TFM)

Gels of 5 kPa were fabricated with embedded fluorescent beads to conduct traction force microscopy (TFM). Briefly, to make a single layer of fluorescent bead (Fluka, 1 μm rhodamine), beads (1:50) were added to the pre-polymer solution (25 μl) and solidified over the normal gel of 5 kPa. Gel was then functionalized as described above. Cells were seeded on the gels, after 24hr of cell seeding, images of stressed (before lysing) and unstressed gel (after lysing with 1% triton-X) were captured by the EVOS FL Auto (Invitrogen). An average of 20 cells were analyzed per conditions. A MATLAB algorithm was used to determine the cell-generated displacement field and traction forces as previously described (5).

### Statistical analysis

Data is presented as means ± standard error of the mean. Unpaired student t-test was used for two sample and one-way ANOVA for three samples. All these test were performed assuming non-gaussian distribution of the sample values. Statistical significant differences were defined as: * =p < 0.05 otherwise stated.

## References

1. Pittenger MF, et al. (1999) Multilineage potential of adult human mesenchymal stem cells. Science 284(5411):143–147.

2. Ranganath SH, Levy O, Inamdar MS, Karp JM (2012) Harnessing the mesenchymal stem cell secretome for the treatment of cardiovascular disease. Cell stem cell 10(3):244–258.

3. Tuan RS (2013) The coming of age of musculoskeletal tissue engineering and regeneration. Nat Rev Rheumatol 9(2):74.

4. Wang LT, et al. (2016) Human mesenchymal stem cells (MSCs) for treatment towards immune- and inflammation-mediated diseases: review of current clinical trials. J Biomed Sci 23(1):76.

5. Caplan AI, Bruder SP (2001) Mesenchymal stem cells: building blocks for molecular medicine in the 21st century. Trends Mol Med 7(6):259–264.

6. Friedenstein A, Latzinik N, Grosheva A, Gorskaya U (1982) Marrow microenvironment transfer by heterotopic transplantation of freshly isolated and cultured cells in porous sponges. Exp Hematol 10(2):217–227.

7. Ren G, et al. (2012) Concise Review: Mesenchymal Stem Cells and Translational Medicine: Emerging Issues. Stem Cells Transl Med 1(1):51–58.

8. Wagner W, et al. (2008) Replicative senescence of mesenchymal stem cells: A continuous and organized process. PloS One 3(5):e2213.

9. Banfi A, et al. (2000) Proliferation kinetics and differentiation potential of ex vivo expanded human bone marrow stromal cells: Implications for their use in cell therapy. Exp Hematol 28(6):707–715.

10. Bonab MM, et al. (2006) Aging of mesenchymal stem cell in vitro. BMC Cell Biol 7(1):14.

11. Wagner W, Ho AD, Zenke M (2010) Different Facets of Aging in Human Mesenchymal Stem Cells. Tissue Eng Part B Rev 16(4):445–453.

12. Honczarenko M, et al. (2006) Human Bone Marrow Stromal Cells Express a Distinct Set of Biologically Functional Chemokine Receptors. Stem cells 24(4):1030–1041.

13. De Becker A, Van Riet I (2016) Homing and migration of mesenchymal stromal cells: How to improve the efficacy of cell therapy? World J Stem Cells 8(3):73.

14. Kassem M (2006) Stem cells: Potential therapy for age-related diseases. Ann N Y Acad Sci 1067(1):436–442.

15. Ullah I, Subbarao RB, Rho GJ (2015) Human mesenchymal stem cells-current trends and future prospective. Biosci Rep 35(2):e00191.

16. Saei Arezoumand K, Alizadeh E, Pilehvar-Soltanahmadi Y, Esmaeillou M, Zarghami N (2017) An overview on different strategies for the stemness maintenance of MSCs. Artif Cells, Nanomedicine Biotechnol 45(7):1255–1271.

17. NIH Stem Cell Information Home Page. In Stem Cell Information [World Wide Web site]. Bethesda, MD: National Institutes of Health, U.S. Department of Health and Human Services, 2016 [cited July 3, 2018] Available at <//stemcells.nih.gov/info/basics/4.htm>

18. Anderson HJ, Sahoo JK, Ulijn RV, Dalby MJ (2016) Mesenchymal Stem Cell Fate: Applying Biomaterials for Control of Stem Cell Behavior. Front Bioeng Biotechnol 4:38.

19. Murphy WL, McDevitt TC, Engler AJ (2014) Materials as stem cell regulators. Nat Mater 13(6):547.

20. Lutolf MP, Gilbert PM, Blau HM (2009) Designing materials to direct stem-cell fate. Nature 462(7272):433–441.

21. Engler AJ, Sen S, Sweeney HL, Discher DE (2006) Matrix Elasticity Directs Stem Cell Lineage Specification. Cell 126(4):677–689.

22. Yeung T, et al. (2005) Effects of substrate stiffness on cell morphology, cytoskeletal structure, and adhesion. Cell Motil Cytoskeleton 60(1):24–34.

23. Gilbert PM, et al. (2010) Substrate elasticity regulates skeletal muscle stem cell self-renewal in culture. Science 329(5995):1078–1081.

24. Winer JP, Janmey PA, McCormick ME, Funaki M (2008) Bone Marrow-Derived Human Mesenchymal Stem Cells Become Quiescent on Soft Substrates but Remain Responsive to Chemical or Mechanical Stimuli. Tissue Eng Part A 15(1):147–154.

25. Zhang D, Kilian KA (2013) The effect of mesenchymal stem cell shape on the maintenance of multipotency. Biomaterials 34(16):3962–3969.

26. Lee J, Abdeen AA, Kim AS, Kilian KA (2015) Influence of Biophysical Parameters on Maintaining the Mesenchymal Stem Cell Phenotype. ACS Biomater Sci Eng 1(4):218–226.

27. Cesarz Z, Tamama K (2016) Spheroid Culture of Mesenchymal Stem Cells. Stem Cells Int 2016:9176357

28. McMurray RJ, et al. (2011) Nanoscale surfaces for the long-term maintenance of mesenchymal stem cell phenotype and multipotency. Nat Mater 10(8):637–644.

29. Binato R, et al. (2013) Stability of human mesenchymal stem cells during in vitro culture: Considerations for cell therapy. Cell Prolif 46(1):10–22.

30. Jalali S, Tafazzoli-Shadpour M, Haghighipour N, Omidvar R, Safshekan F (2015) Regulation of Endothelial Cell Adherence and Elastic Modulus by Substrate Stiffness. Cell Commun Adhes 22(2–6):79–89.

31. Tee SY, Fu J, Chen CS, Janmey PA (2011) Cell shape and substrate rigidity both regulate cell stiffness. Biophys J 100(5):L25–L27.

32. Rumman M, et al. (2018) Induction of quiescence (G0) in bone marrow stromal stem cells enhances their stem cell characteristics. Stem Cell Res 30:69–80.

33. Nishio K, Inoue A, Qiao S, Kondo H, Mimura A (2001) Senescence and cytoskeleton: Overproduction of vimentin induces senescent-like morphology in human fibroblasts. Histochem Cell Biol 116(4):321–327.

34. Frescas D, et al. (2017) Senescent cells expose and secrete an oxidized form of membrane-bound vimentin as revealed by a natural polyreactive antibody. Proc Natl Acad Sci 114(9):E1668–E1677.

35. Kundrotas G, et al. (2016) Identity, proliferation capacity, genomic stability and novel senescence markers of mesenchymal stem cells isolated from low volume of human bone marrow. Oncotarget 7(10):10788–10802.

36. Vidal MA, Walker NJ, Napoli E, Borjesson DL (2011) Evaluation of senescence in mesenchymal stem cells isolated from equine bone marrow, adipose tissue, and umbilical cord tissue. Stem cells and development 21(2):273–283.

37. Meng X, et al. (2017) MicroRNA profiling analysis revealed different cellular senescence mechanisms in human mesenchymal stem cells derived from different origin. Genomics 109(3):147–157.

38. Chowdhury F, et al. (2010) Soft substrates promote homogeneous self-renewal of embryonic stem cells via downregulating cell-matrix tractions. PloS One 5(12):e15655.

39. McBeath R, Pirone DM, Nelson CM, Bhadriraju K, Chen CS (2004) Cell shape, cytoskeletal tension, and RhoA regulate stem cell lineage commitment. Dev Cell 6(4):483–495.

40. McGrail DJ, McAndrews KM, Dawson MR (2013) Biomechanical analysis predicts decreased human mesenchymal stem cell function before molecular differences. Exp Cell Res 319(5):684–696.

41. Wall ME, Bernacki SH, Loboa EG (2007) Effects of Serial Passaging on the Adipogenic and Osteogenic Differentiation Potential of Adipose-Derived Human Mesenchymal Stem Cells. Tissue Eng 13(6):1291–1298.

42. Neuhuber B, Swanger SA, Howard L, Mackay A, Fischer I (2008) Effects of plating density and culture time on bone marrow stromal cell characteristics. Exp Hematol 36(9):1176–1185.

43. Kilian KA, Bugarija B, Lahn BT, Mrksich M (2010) Geometric cues for directing the differentiation of mesenchymal stem cells. Proc Natl Acad Sci 107(11):4872–4877.

44. Basciano L, et al. (2011) Long term culture of mesenchymal stem cells in hypoxia promotes a genetic program maintaining their undifferentiated and multipotent status. BMC Cell Biol 12(1):12.

45. Freund A, Laberge RM, Demaria M, Campisi J (2012) Lamin B1 loss is a senescence-associated biomarker. Mol Biol Cell 23(11):2066–2075.

46. Bellotti C, et al. (2016) Detection of mesenchymal stem cells senescence by prelamin A accumulation at the nuclear level. Springerplus 5(1):1427.

## References

1. Pelham RJ, Wang Yl (1997) Cell locomotion and focal adhesions are regulated by substrate flexibility. Proc Natl Acad Sci 94(25):13661–13665.

2. Tse JR, Engler AJ (2010) Preparation of hydrogel substrates with tunable mechanical properties. Curr Protoc Cell Biol 47(SUPPL. 47):10.16.1–10.16.16.

3. Venugopal B, Mogha P, Dhawan J, Majumder A (2018) Cell density overrides the effect of substrate stiffness on human mesenchymal stem cells’ morphology and proliferation. Biomater Sci 6(5):1109–1119.

4. Cristofalo VJ, Allen RG, Pignolo RJ, Martin BG, Beck JC (1998) Relationship between donor age and the replicative lifespan of human cells in culture: A reevaluation. Proc Natl Acad Sci 95(18):10614–10619.

5. Butler JP, Tolic-Norrelykke IM, Fabry B, Fredberg JJ (2002) Traction fields, moments, and strain energy that cells exert on their surroundings. AJP Cell Physiol 282(3):C595–C605.

